# Rod-genesis driven by *mafba* in an *nrl* knockout zebrafish model with altered photoreceptor compositions and progressive retinal degeneration

**DOI:** 10.1101/2021.09.29.462301

**Authors:** Fei Liu, Yayun Qin, Yuwen Huang, Pan Gao, Jingzhen Li, Shanshan Yu, Danna Jia, Xiang Chen, Yuexia Lv, Jiayi Tu, Kui Sun, Yunqiao Han, James Reilly, Xinhua Shu, Qunwei Lu, Zhaohui Tang, Chengqi Xu, Daji Luo, Mugen Liu

## Abstract

The neural retina leucine zipper (*NRL*) is an essential gene for the fate determination and differentiation of rod photoreceptors in mammals. Mutations in *NRL* have been associated with autosomal recessive enhanced S-cone syndrome and autosomal dominant retinitis pigmentosa. However, the exact role of Nrl in regulating the development and maintenance of photoreceptors in zebrafish, a popular animal model used for retinal degeneration and regeneration studies, has not been fully determined. In this study, we generated an *nrl* knockout zebrafish model by CRISPR-Cas9 technology and observed a surprising phenotype characterized by the reduction but not total elimination of rods and the over-grown of green-cones. By tracing the developmental process of rods, we discovered two waves of rod genesis in zebrafish, emerging at the embryonic stage with an *nrl*-dependent pattern and the post-embryonic stage with an *nrl*-independent pattern, respectively. Through bulk and single-cell RNA sequencing, we constructed the gene expression profiles for the whole retinal tissues and each of the retinal cell types in WT and *nrl* knockout zebrafish. We detected the rod/green-cone intermediate photoreceptors in *nrl* knockout zebrafish, suggesting that there may be a kind of rod/green-cone bipotent precursors and its fate choice between rod and green-cone is controlled by *nrl*. Besides, we identified the *mafba* gene as a novel regulator for *nrl*-independent rods, based on the cell-type-specific expression pattern and the retinal phenotype of *nrl*/*mafba* double knockout zebrafish. Furthermore, the altered photoreceptor compositions and abnormal gene expression caused progressive retinal degeneration and subsequent regeneration in *nrl* knockout zebrafish. Our work revealed a novel function of *mafba* gene in rod development and established a more suitable model for the developmental processes and regulatory mechanisms of rod and green-cone photoreceptors in zebrafish.

**Author Summary:** Vision is mediated by two types of light-sensing cells named rod and cone photoreceptors in animal eyes. Abnormal generation, dysfunction or death of photoreceptor cells can all cause irreversible vision problems. *NRL* is the most important gene for the occurrence and function of rod cells in mice and humans. Surprisingly, we found that in zebrafish, a popular animal model used for mimicking human diseases and testing treatments, there are two types of rod cells and breaking the function of *nrl* gene only affects the generation of rod cells at the embryonic stage but not the juvenile and adult stages. The rod cells produced later are proved to be driven by the *mafba* gene, which has not been reported to play a role in rod cells. In addition to the reduction of rod cells, deletion of *nrl* also results in the occurrence of rod/green-cone hybrid cells and the increasing number of green-cones. The combination of changes at cellular and molecular levels finally lead to a disease condition named retinal degeneration. These findings add new knowledge to the research field and highlight the conserved and species-specific regulatory mechanisms of photoreceptor development and maintenance.

## Introduction

Vision is involved in many fundamental behaviors of animals such as navigation, foraging, predator avoidance, and mate choice [1]. In human beings, inherited retinal diseases are a major cause of irreversible vision impairment and blindness, resulting in serious physical discomfort, mental problems, and economic burden for patients [2,3]. There are two types of light-sensing cells named rod and cone photoreceptors in vertebrate eyes. Rods are highly sensitive to dim light and function under conditions of low light, while cones respond to bright light and mediate color vision [4,5]. Rods and cones are derived from the same progenitor cells regulated by several extrinsic signals and intrinsic transcription factors such as *CRX, NRL, NR2E3* [6,7]. Exploring the fundamental mechanisms controlling the fate determination and differentiation of photoreceptors in diverse animal models is of great value for understanding the development and evolution of photoreceptors, and also the pathogenesis of retinal degenerative diseases.

*NRL* (neural retina leucine zipper), a large MAF family transcription factor expressed in the retina and pineal gland [8,9], has been identified as the master gene determining the cell fate of rod versus cone photoreceptors in mammals. Disruption of *Nrl* in mice results in the complete loss of rod photoreceptors, accompanied by a large excess of S-cone-like photoreceptors [10,11]. Similarly, loss-of-function mutations in *NRL* cause autosomal recessive enhanced S-cone syndrome (ESCS) in humans, which is characterized by the absence of rod response and enhanced S-cone function [12-14]. Moreover, ectopic expression of *Nrl* transforms the fate of cone precursors to rods in mice [15], suggesting that *NRL* is essential and sufficient for the fate determination and differentiation of rod photoreceptors. Additionally, NRL activates the expression of many rod-specific genes (such as *RHO, GNAT1, PDE6A, REEP6, MEF2C*) synergistically with CRX and NR2E3 [16-21], and represses the cone-specific genes (such as *Thrb* and *Opn1sw*) either directly or by activating the expression of *NR2E3* [22,23].

In recent years, zebrafish has emerged as a useful animal model for retinal degeneration and regeneration studies [24-27]. Interestingly, increasing evidence suggests that the current working model of photoreceptor development, which is largely based on studies in mice and humans, may not perfectly match the observations in zebrafish [28-31]. For example, a portion of rods are derived from S-cone precursors in mice, but this phenomenon was not observed in zebrafish [29]. In addition, our recent study showed that deletion of *nr2e3*, another key determining factor of rod fate [32,33], eliminates rods completely but doesn’t increase the number of UV- or blue-cones (corresponding to S-cones) in zebrafish [30], which disagrees with the phenotypes of *Nr2e3* or *Nrl* knockout mice [10,32]. Whether rods could be derived from S-cone or other cone precursors in zebrafish and how this process is regulated remain unclear.

In this study, we generated an *nrl* knockout zebrafish model by CRISPR-Cas9 technology and systematically investigated the developmental processes of rod photoreceptors. Surprisingly, we found two waves of rod genesis in zebrafish, which could be distinguished as *nrl*-dependent at embryonic stage and *nrl*-independent at post-embryonic stage. In addition, we observed the continuous increase of green-cones in adult *nrl* knockout zebrafish. The cellular and molecular mechanisms underlying this unexpected retinal phenotype were further investigated, and a modified model for the developmental processes and regulatory mechanisms of rod and green-cone photoreceptors in zebrafish was established.

## Results

### Knockout of *nrl* leads to reduction but not complete loss of rods in zebrafish

The *nrl* knockout zebrafish was generated by CRISPR-Cas9 targeting the exon 2 of *nrl*, which encodes the conserved N-terminal minimal transactivation domain (Fig 1A). Through several rounds of crossing and mutation screening, we obtained a homozygous *nrl* mutant zebrafish line carrying a frameshift mutation (c.230_237del8, p.P77Hfs*4) (Fig 1B). The mutation was predicted to cause a severely truncated form of Nrl protein without intact functional domains. The *nrl* mRNA levels were significantly up-regulated in this mutant zebrafish line (named as *nrl*-KO hereinafter) (Fig 1C), possibly due to a negative feedback mechanism [34]. No alternative transcript of *nrl* was detected in zebrafish retinas. We have also tried to detect the protein levels of Nrl in WT and *nrl*-KO retinas. Unfortunately, our customized antibodies against zebrafish Nrl could not work in western blot and immunofluorescence assays.

**Fig 1.**
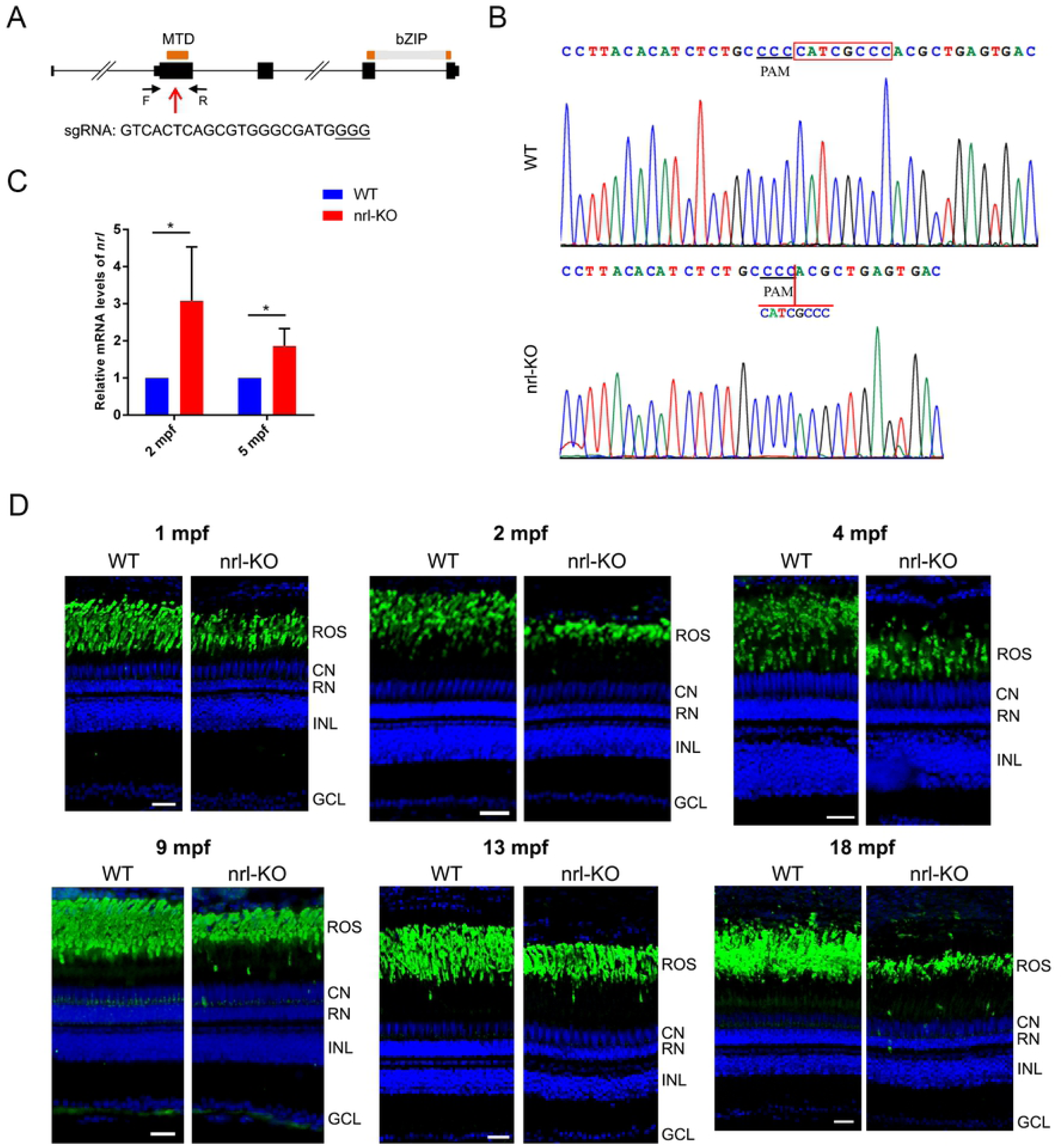
Knockout of *nrl* leads to reduction but not complete loss of rods in zebrafish. (A) The gene structure of zebrafish *nrl* and the CRISPR-Cas9 target site used for *nrl* knockout. Orange boxes, the exons encoding the MTD and bZIP domains; red arrow, the CRISPR-Cas9 target site; black arrows, primers used for mutation detection. (B) Sequencing validation of the homozygous *nrl* del8 mutation (c.230_237del8). (C) The *nrl* mRNA levels were detected by qPCR in WT and *nrl*-KO zebrafish. The data are shown as mean with SD (n=3). *, p < 0.05. (D) Immunostaining of rod outer segments using the anti-Rho antibody on retinal sections of WT and *nrl*-KO zebrafish from 1 mpf to 18 mpf. Scale bars: 25 μm. ROS, rod outer segment; CN, cone nuclear layer; RN, rod nuclear layer; INL, inner nuclear layer; GCL: ganglion cell layer.

Histological analysis was performed to examine the retinal morphology of *nrl*-KO zebrafish. The thicknesses of photoreceptor and outer segment layers were markedly reduced in *nrl*-KO retinas (S1 Fig). The rosette-like anomalies, which are commonly observed in the outer nuclear layer (ONL) of *Nrl* and *Nr2e3* knockout mice [35,36], were not observed in *nrl*-KO zebrafish (S1A and S1D Fig). Surprisingly, the presence of rod nuclei (labelled with RN in S1B and S1E Fig) suggested that rods might not be completely eliminated by *nrl* knockout in zebrafish.

Deletion of *Nrl* has been reported to cause complete loss of rods in mice [10,11]. To more clearly show whether there are rods in *nrl*-KO zebrafish, the anti-zebrafish Rho antibody was used to specifically label rods through immunostaining. Indeed, we observed rod outer segments in the retinal sections of *nrl*-KO zebrafish from 14 dpf (day post-fertilization) to 18 mpf (month post-fertilization) (Fig 1D and S2 Fig). The thicknesses of rod outer segment layer and the number of rod nuclei were both significantly reduced when compared with WT groups. The density of rod outer segments was also significantly reduced in *nrl*-KO zebrafish, especially in the ventral retina, as shown by the retinal whole-mount immunostaining (S3 Fig). Interestingly, we noticed that rods could not be generated in the marginal regions of *nrl*-KO retinas at 14 dpf and 1 mpf (S2A and S2B Fig). However, at the age of 2 mpf and 4 mpf, these regions were covered by rods as in other retinal regions (S2C and S2D Fig), suggesting a central-to-peripheral developmental pattern of rods in *nrl*-KO zebrafish. Our results indicated that knockout of *nrl* in zebrafish reduced the number of rods rather than completely eliminating them, which was quite different from the retinal phenotypes of *Nrl* and *Nr2e3* knockout mice [10,37].

### Identification of two waves of rod genesis in zebrafish

To trace the developmental processes of rods in *nrl*-KO zebrafish, we used the Tg(rho:EGFP) transgene zebrafish to label rods with EGFP (enhanced green fluorescent protein) *in vivo*. In WT zebrafish, rods first appeared in the retinal ventral edge and then spread to the entire retina with rapidly increased fluorescence intensity (Fig 2A-2C). However, we didn’t observe the fluorescence of rods until 7 dpf in *nrl*-KO zebrafish (Fig 2A). Immunostaining of another rod-specific protein Gnat1 on retinal sections also showed the complete loss of rods in *nrl*-KO retinas at 7 dpf (S4A Fig). Interestingly, after 10 dpf, the fluorescence of rods in *nrl*-KO zebrafish became obvious and increased with age rapidly (Fig 2B). The distribution of rods in *nrl*-KO retinas became more and more extensive as revealed by the flattened retinal whole-mounts from 1 mpf to 3 mpf, although the expression levels of rod opsin (reflected by fluorescence intensities) were still lower than WT controls (Fig 2C).

**Fig 2.**
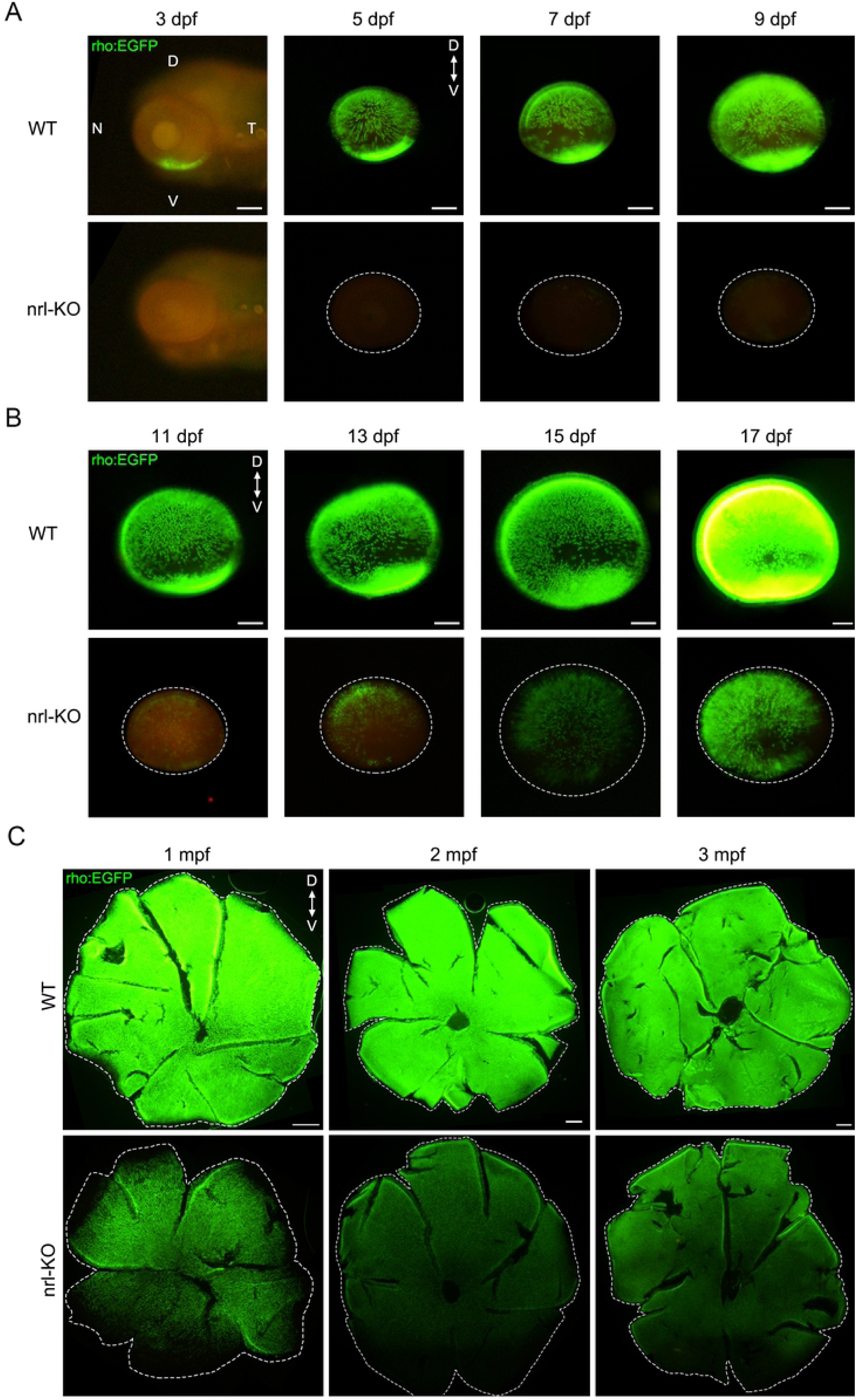
Tracing the developmental processes of rods in WT and *nrl*-KO zebrafish. Rods were labeled with EGFP by crossing WT and *nrl*-KO zebrafish with the Tg(rho:EGFP) transgene zebrafish. Fluorescence were observed every 2 days from 3 dpf. The representative images of WT and *nrl*-KO retinas are shown in (A) for 3-9 dpf and (B) for 11-17 dpf. The dotted circles indicate the boundaries of retinas. D, dorsal; V, ventral; N, nasal; T, temporal. Scale bars: 100 μm. (C) Representative images of flattened whole-mount retinas from WT and *nrl*-KO zebrafish at 1-3 mpf. The dashed lines indicate the edges of the retinas. The distribution of rods in WT and *nrl*-KO retinas are shown. Scale bars: 200 μm.

To show the EGFP-labeled rods more clearly, retinal sections of Tg(rho:EGFP) transgene zebrafish were prepared and observed at high magnification (S4B Fig). The rod numbers were obviously reduced in *nrl*-KO retinas, and the fluorescence of every single rod was also significantly lower than WT zebrafish. Besides, rods were absent in the marginal regions of *nrl*-KO retina at 1 mpf, which was consistent with our immunostaining results shown above (S2A and S2B Fig). Taken together, these results suggested that there are two waves of rod genesis in zebrafish. Knockout of *nrl* eliminates the first wave of rod genesis starting at about 3 dpf (embryonic stage), but doesn’t affect the second wave of rod genesis starting at 7-10 dpf (juvenile stage), which were named as the *nrl*-dependent and *nrl*-independent rods, respectively.

### Decreased expression of rod-specific genes and increased green-cones in adult *nrl*-KO zebrafish

In order to investigate the transcriptome dynamics of *nrl*-KO retinas, we performed RNA-seq analysis using 2 mpf WT and *nrl*-KO retinas. A total of 386 differentially expressed genes (fold change ≥2, adjusted p-value ≤0.05, and FPKM ≥1) were detected (Fig 3A and S1 Dataset). The functions and related biological processes of the top 100 differentially expressed genes were classified into several categories (S2 Dataset). Interestingly, we found that the most affected functional categories were inflammatory and immune response, signal transduction, metabolic enzymes, cell adhesion and extracellular matrix, transport, cell protection and death (Fig 3B). These altered biological functions were similar to those identified in *Nrl* knockout mice[38], which might reflect the transient degeneration and retinal remodeling in *nrl*-KO zebrafish.

**Fig 3.**
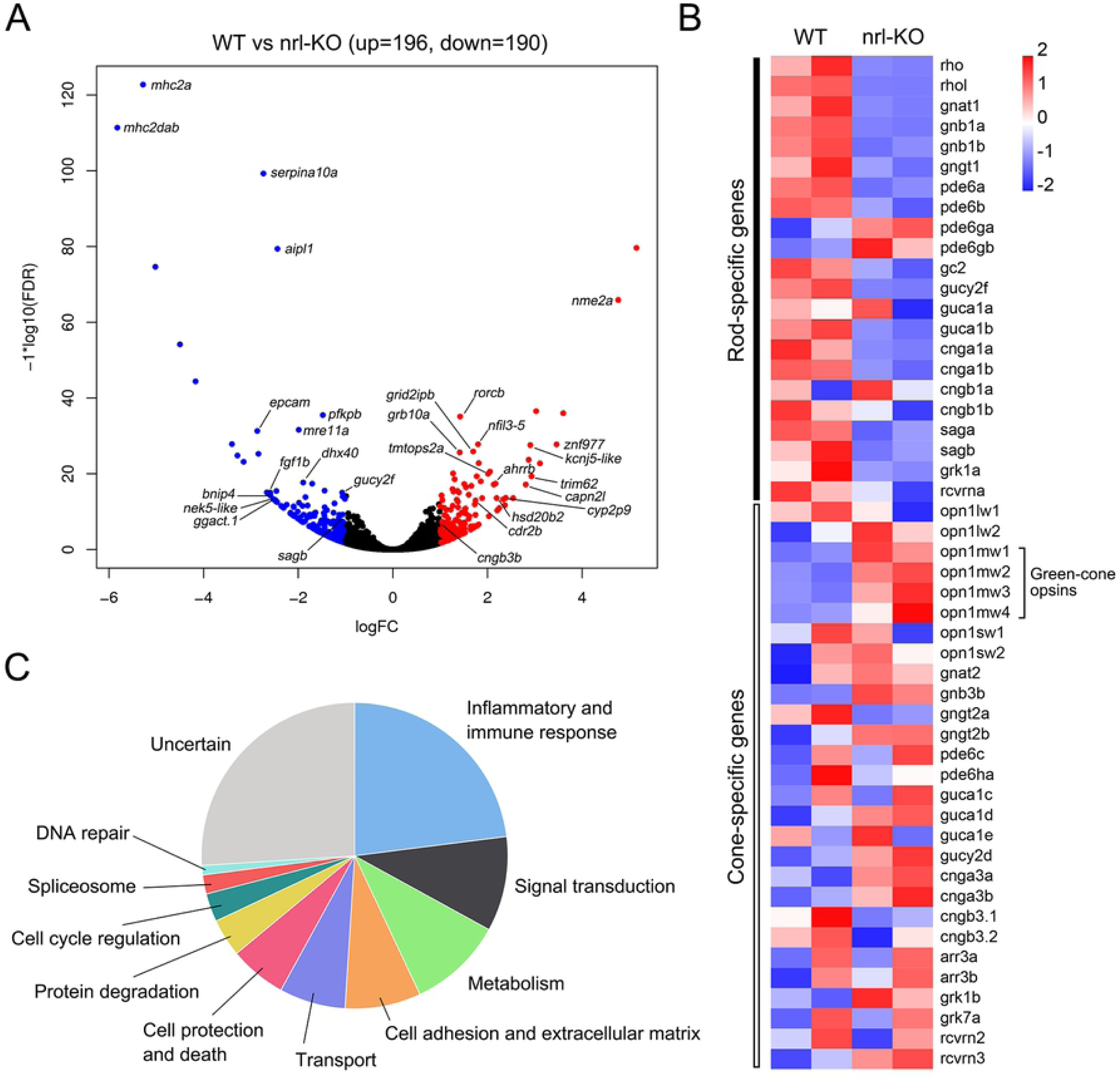
Differentially expressed genes between WT and *nrl*-KO retinas detected by RNA-seq. (A) The volcano plot shows the 386 differentially expressed genes (196 up-regulated and 190 down-regulated) between WT and *nrl*-KO retinas at 2 mpf. The red and blue points indicate the up-regulated and down-regulated genes, respectively. (B) Functional categories of the top 100 differentially expressed genes. (C) Expression patterns of rod- and cone-specific phototransduction genes in *nrl*-KO retinas shown by the heatmap.

To our surprise, only few differentially expressed genes (*aipl1, nxnl1, sagb, gucy2f, rpgra*, and *cnga3b*) were closely related to the development or degeneration of retinas. Because we have observed the reduction of rods in *nrl*-KO zebrafish, we directly examined the expression of rod- and cone-specific phototransduction genes based on our RNA-seq data. As expected, most of the rod-specific genes were down-regulated (adjusted p-value ≤0.05) in the *nrl*-KO group (Fig 3C). However, the fold changes were mostly less than 2 folds, which were ignored by the routine bioinformatic pipelines. Meanwhile, the mRNA levels of cone-specific genes were not significantly changed, except for the obviously up-regulated green-cone opsins (Fig 3C).

To confirm the RNA-seq results, we detected the protein levels of representative rod-specific (*rho, gnat1*, and *gnb1*) and cone-specific (*gnat2* and *gnb3*) genes from 14 dpf to 3 mpf by western blot. The rod-specific genes were significantly down-regulated in *nrl*-KO retinas (Fig 4A and 4B). Meanwhile, the protein levels of cone-specific genes were unchanged before 2 mpf, but significantly increased at 3 mpf (Fig 4A and 4B). We also performed qPCR to detect the mRNA levels of cone opsin genes at 2 and 5 mpf. The green-cone opsins (*opn1mw1, opn1mw2, opn1mw3, opn1mw4*) were significantly up-regulated at both time points, while the expression of UV-, blue-, and red-cone opsins were almost unchanged (Fig 4C). These results correlated well with our RNA-seq data and suggested that the reduction of rod-specific genes and increase of cone-specific genes (more particularly green-cone genes) may co-exist in *nrl*-KO zebrafish.

**Fig 4.**
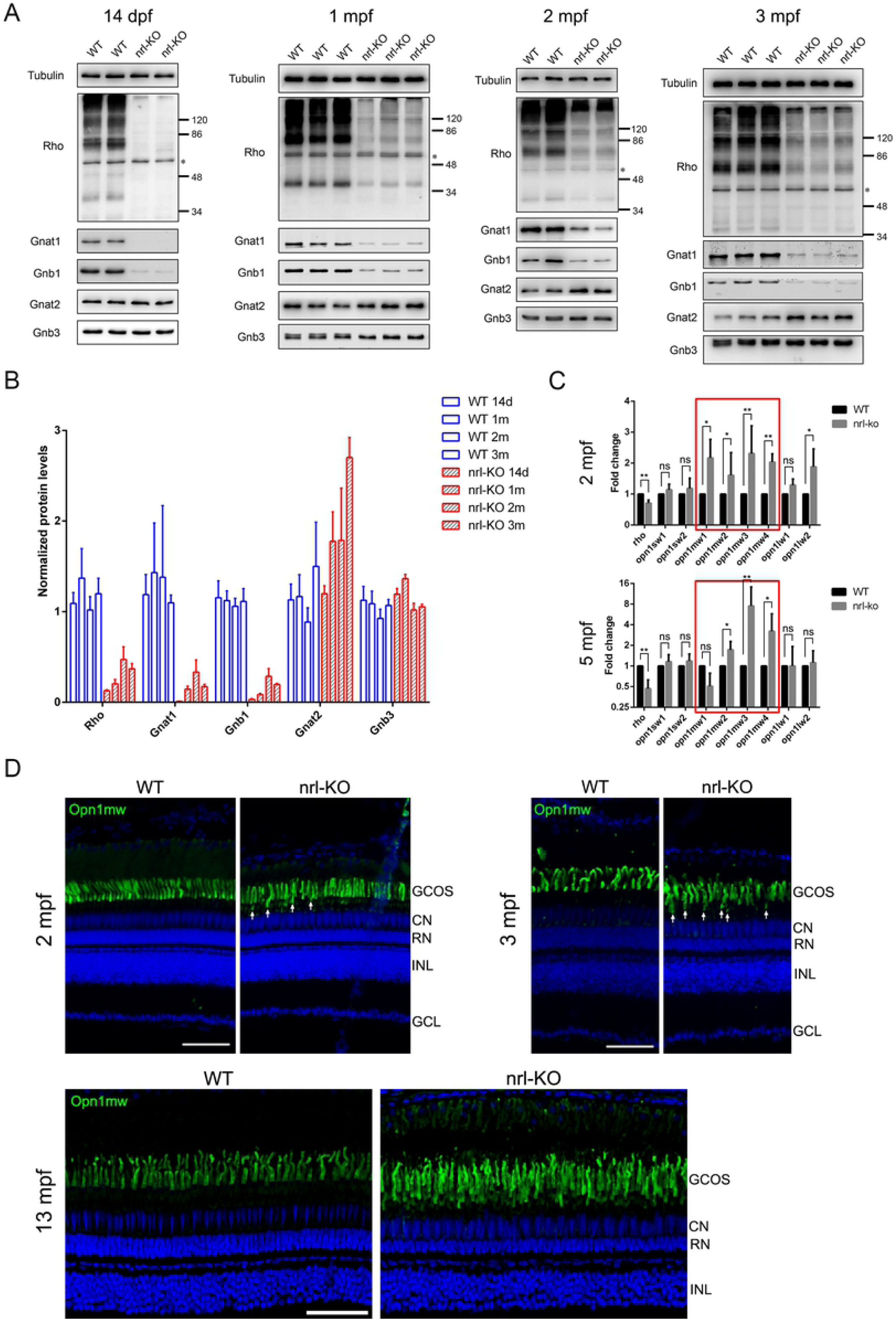
Decreased expression of rod-specific genes and increased green-cones in adult *nrl*-KO zebrafish. (A) The protein levels of rod-specific (*rho, gnat1*, and *gnb1*) and cone-specific (*gnat2* and *gnb3*) genes in WT and *nrl*-KO retinas from 14 dpf to 3 mpf were detected by western blot. Tubulin was served as the loading control. The asterisk indicates a non-specific band. (B) Quantitative results of the normalized protein levels from at least three independent experiments. The data are shown as mean with SD. (C) The mRNA levels of rod and cone opsins were detected by qPCR in WT and *nrl*-KO retinas at 2 and 5 mpf. The data are shown as mean with SD (n=3). ns, non-significant; *, p < 0.05; **, p < 0.01. (D) Detection of green-cones on retinal sections of WT and *nrl*-KO zebrafish at 2, 3, and 13 mpf by immunostaining using the anti-Opn1mw antibody. The dorsal retinal regions are shown. White arrows indicate the mislocated green-cone outer segments. GCOS, outer segments of green-cones; CN, cone nuclear layer; RN, rod nuclear layer; INL, inner nuclear layer; GCL, ganglion cell layer. Scale bars: 50 μm.

To validate whether the number of green-cones is also increased, we performed immunostaining using the anti-Opn1mw (zebrafish green-cone opsin) antibody on retinal sections of WT and *nrl*-KO zebrafish. No obvious changes were observed in *nrl*-KO retinas at 2 mpf and 3 mpf, except for some newly formed green-cone outer segments located below the normal single-row arrangement (Fig 4D). However, at 13 mpf, 2-3 rows of green-cone outer segments could be observed, accompanied by the remarkable increase of cone nuclei (Fig 4D), suggesting a remarkable increase of green-cones. The expression levels of genes involved in cone development (such as *rx1, tbx2a, tbx2b, six7*, and *thrb*) were not significantly affected as suggested by the RNA-seq data, indicating that the over-growth of green-cones since 3 mpf might be medicated by an unknown mechanism.

### Single-cell RNA-seq revealed the cell compositions and rod/green-cone intermediate photoreceptors in *nrl* knockout retinas

The retinal cell compositions and cell-type-specific gene expression patterns were determined by single-cell RNA-seq (scRNA-seq) analysis using WT and *nrl*-KO zebrafish at 5 mpf. After filtering out the invalid cells, 10374 cells (4682 from WT group and 5692 from nrl-KO group) were classified into 24 clusters by unsupervised cell clustering analysis (Fig 5A). The relative mRNA levels of all genes expressed in the 24 cell types were determined (S3 Dataset). According to the expression patterns of known marker genes for each retinal cell type [39-42], the 24 cell clusters were identified as rods (3 subtypes), cones (2 subtypes), horizontal cells, bipolar cells (12 subtypes), amacrine cells, RGCs, RPE, Muller glia, astrocytes, and microglia (S5 Fig, S4 Dataset).

**Fig 5.**
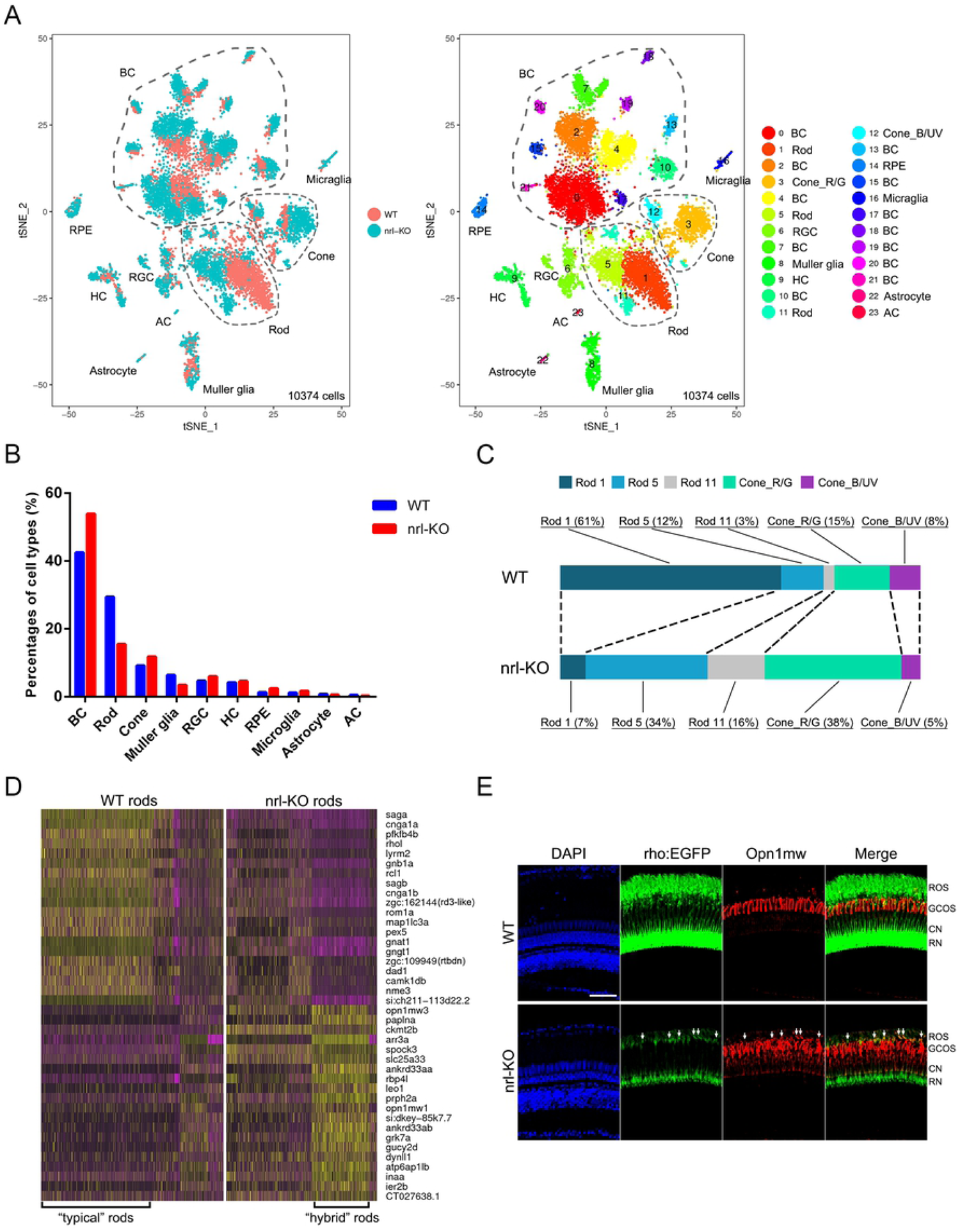
Single-cell RNA-seq analysis in WT and *nrl*-KO retinas. **(A)** tSNE visualization of the unsupervised cell clusters from 5-month-old WT and *nrl*-KO zebrafish. Left, the distribution of cell clusters between WT and *nrl*-KO groups. Right, the retinal cell types identified by scRNA-seq (see also Fig. S5). BC, bipolar cells; Cone_R/G, red and green cones; Cone_B/UV, UV and blue cones; RGC, retinal ganglion cells; HC, horizontal cells; AC, amacrine cells; RPE, retinal pigment epitheliums. (B) The proportions of each cell type in WT and *nrl*-KO retinas. (C) The cell compositions and proportions of photoreceptor sub-clusters in WT and *nrl*-KO retinas. (D) The heatmap shows the clustering result of the top 20 down-regulated and top 20 up-regulated genes in WT and *nrl*-KO rods. Yellow, high expression; purple, low expression. (E) Mis-expression of the green-cone opsin in a proportion of rods in *nrl*-KO zebrafish. The dorsal retinal regions of WT and *nrl*-KO zebrafish at 3 mpf are shown. Green, EGFP-labeled rods. Red, immunofluorescence signals of the anti-Opn1mw antibody. ROS, outer segments of rods; GCOS, outer segments of green cones; CN, cone nuclear layer; RN, rod nuclear layer. Scale bar: 50 μm.

The percentages of all cell types in WT and *nrl*-KO retinas were analyzed (Fig 5B). In *nrl*-KO zebrafish, the percentage of rods was reduced by about 50%, while cones (more accurately the red/green cones) showed an obvious increase (Fig 5B and 5C). As for the three rod subtypes (rod-1, rod-5, and rod-11), rod-1 was the major rod subtype in WT zebrafish (80% of all rods), which was reduced to 13% in *nrl*-KO zebrafish. Conversely, rod-5 and rod-11 were significantly increased in *nrl*-KO zebrafish (34% and 16% of all rods). These data suggested that knockout of *nrl* may affect both the fate determination of rods versus green-cones and the ratio of rod subtypes.

The gene expression signatures for the three rod subtypes were constructed based on the gene expression profiles through unsupervised clustering analysis. The top 40 discriminative genes between WT and *nrl*-KO rods are shown in the clustering heatmap (Fig 5D). The leftmost cells showed high expression of rod-specific genes (such as *rho, rhol, saga, sagb, gnat1*) and very-low expression of cone-specific genes (such as *opn1mw1, opn1mw3, arr3a, grk7a, gucy2d*), representing the “typical” rods in WT zebrafish. By contrast, the rightmost cells showed relatively low levels of rod-specific genes and high levels of cone-specific genes (especially the green-cone opsins), probably representing the “hybrid” rods (rod/green-cone intermediate photoreceptors) in *nrl*-KO zebrafish.

To directly visualize these intermediate photoreceptors, immunostaining was performed on retinal sections of Tg(rho:EGFP) transgene zebrafish using the anti-Opn1mw (green-cone opsin) antibody. The improper expression and accumulation of Opn1mw in some of the rods (labeled by EGFP) were only observed in *nrl*-KO retinas (Fig 5E). These results demonstrated the existence of rod/green-cone intermediate photoreceptors in *nrl*-KO zebrafish.

Previous studies have shown the abnormal chromatin morphology of the S-cone-like cells in *Nrl* or *Nr2e3* knockout mice [10,32]. Interestingly, our scRNA-seq data showed that the *hmgn2* gene, which encodes a non-histone nucleosomal binding protein and contributes to the chromatin plasticity and epigenetic regulation [43], was remarkably up-regulated in *nrl*-KO rods. Through *in situ* hybridization, we found that *hmgn2* was expressed at a very low level in WT rods (S6 Fig). However, in *nrl*-KO zebrafish the expression of *hmgn2* was specifically enhanced in rods (S6 Fig). These results suggested that changes of epigenetic modification might also play a role in the formation of intermediate photoreceptors in *nrl*-KO zebrafish.

### Identification of *mafba* as a novel driving factor for rod development

A reasonable explanation for the occurrence of *nrl*-independent rods is that another gene might at least partly substitute for the functions of *nrl* in driving rod genesis. Genes belonging to the large MAF family (*MAFA, MAFB*, and *MAF*/*c-MAF*) are the most likely candidates due to the high conservation between NRL and them [29,44]. In addition, *MAFA* has been reported to induce the expression of rod-specific genes when ectopically expressed in mice [29,45]. Based on our scRNA-seq data, the gene expression patterns of all large MAF genes and genes involved in photoreceptor development were determined. We found that *mafba*, one of the two *MAFB* orthologous genes in zebrafish, showed the closest relationship with *nrl* and *nr2e3* (Fig 6A). *mafba* was also the only one of the large MAF family genes expressed in all three types of rods with a comparable level to *nrl* and *nr2e3* (S7A Fig).

**Fig 6.**
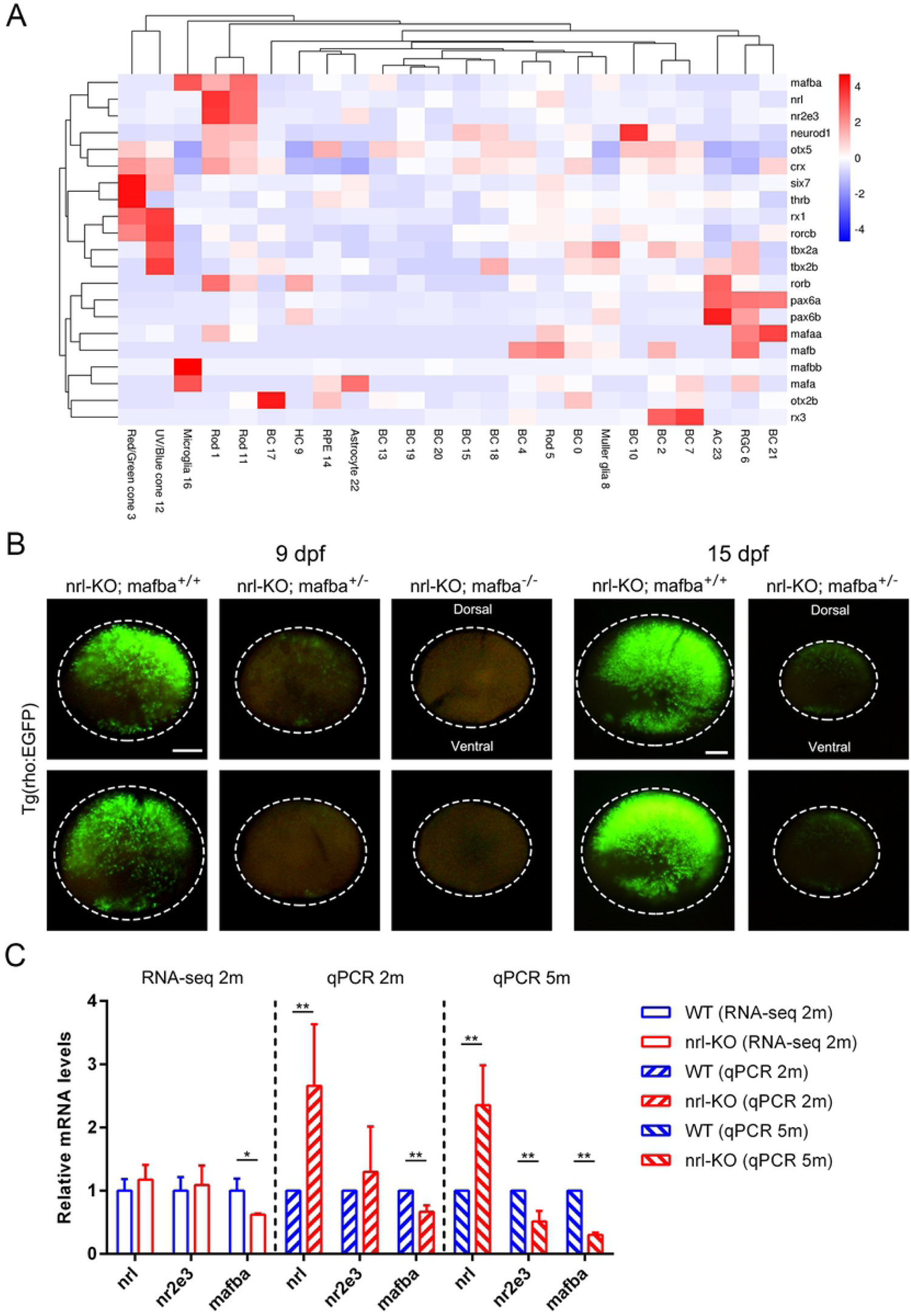
Identification of *mafba* as a novel driving factor for the *nrl*-independent rods. (A) Clustering analysis of the large MAF genes and the genes involved in photoreceptor development using the expression data from scRNA-seq. The cell-type-specific expression pattern of *mafba* was highly similar to those of *nrl* and *nr2e3*. (B) Observing the development of rods (labeled by EGFP) in *nrl*-KO zebrafish with the WT, heterozygous, and homozygous *mafba* mutant allele at 9 dpf (left panel) and 15 dpf (right panel). Knockout of *mafba* additionally reduced the genesis of rods in *nrl*-KO zebrafish. The dotted circles indicate the boundaries of retinas. Scale bars: 100 μm. (C) The expression changes of *nrl, nr2e3*, and *mafba* in *nrl*-KO retinas at 2 mpf and 5 mpf as detected by RNA-seq and qPCR. The data are shown as mean with SD. *, p < 0.05; **, p < 0.01.

To validate whether *mafba* plays a role in rod development, we constructed the *mafba* knockout zebrafish by CRISPR-Cas9 technology (S7B and S7C Fig), and then the *nrl* and *mafba* double knockout zebrafish by crossing the two zebrafish lines. The homozygous *mafba* mutants died at 9-10 dpf with multi-organ developmental defects, as described previously [46,47]. In *nrl*-KO zebrafish, deleting both copies of *mafba* completely eliminated rods at 9 dpf (Fig 6B). Interestingly, deleting one copy of *mafba* also dramatically retarded the development of *nrl*-independent rods at 9 and 15 dpf, suggesting a dose-dependent effect of *mafba* (Fig 6B). We also examined the mRNA levels of *nrl, nr2e3*, and *mafba* through RNA-seq and qPCR. The expression of *mafba* was significantly decreased in *nrl*-KO retinas at 2 and 5 mpf (Fig 6C), excluding the genetic compensation effect caused by *nrl* disruption. Taken together, our results demonstrated that *mafba* is a novel regulatory gene driving rod genesis in zebrafish.

### Progressive retinal degeneration and regeneration in *nrl* knockout zebrafish

The gradual loss of rods is considered as a main cause of the secondary death of cones in retinitis pigmentosa patients [48,49]. As the *nrl*-KO zebrafish showed obvious reduction of rods and excessive increase of green-cones, we wondered whether these developmental abnormalities would lead to retinal degeneration. TUNEL assay was performed to detect cell death on retinal sections from 5 mpf to 13 mpf. Indeed, we observed more apoptotic cells in *nrl*-KO retinas, including rods (predominant), cones, RPE, and inner retinal cells (Fig 7A and 7B). Immunostaining of ZO-1 showed the abnormal morphology of RPE cells in *nrl*-KO retinas (Fig 7C). We also detected the up-regulation of GFAP in aged *nrl*-KO retinas (Fig 7D), which is a hallmark of retinal damage. These results demonstrated the existence of progressive retinal degeneration in *nrl*-KO zebrafish.

**Fig 7.**
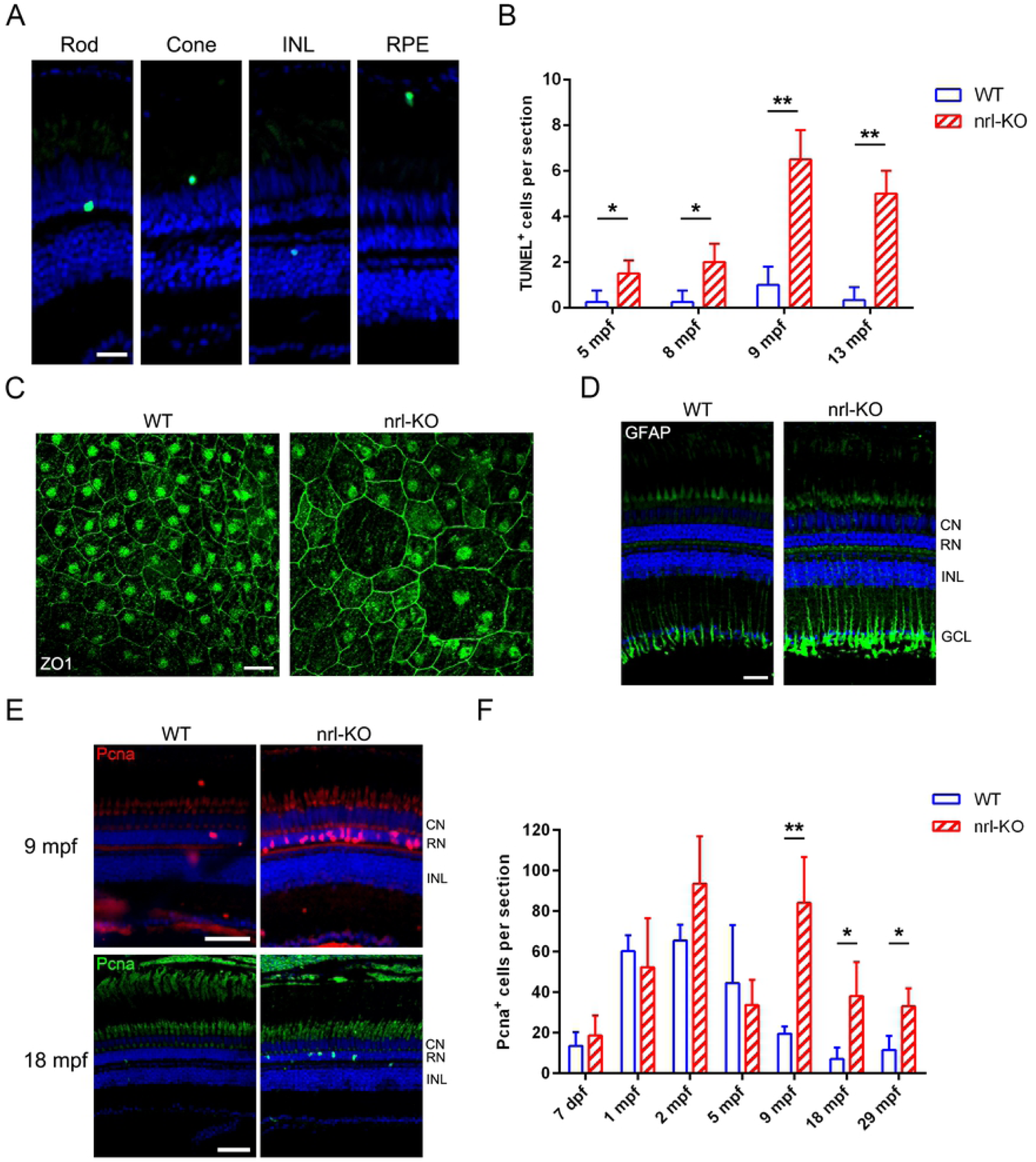
Progressive degeneration and regeneration in *nrl*-KO retinas. **(A)** Apoptosis of rods, cones, RPE and inner retinal cells were detected by TUNEL assays. The representative images are shown. Scale bar: 20 μm. (B) Quantitative results of apoptotic cells per section from 5 mpf to 13 mpf are shown as mean with SD (n=3). *, p < 0.05; **, p < 0.01. (C) Immunostaining of ZO-1 showed the RPE morphology in the retinal whole-mounts of WT and *nrl*-KO zebrafish at 20 mpf. Scale bar: 20 μm. (D) The up-regulation of GFAP in *nrl*-KO retinas detected by immunostaining. CN, cone nuclear layer; RN, rod nuclear layer; INL, inner nuclear layer; GCL, ganglion cell layer. Scale bar: 25 μm. (E) The representative immunostaining images of Pcna, a maker of proliferating cells, showed higher regeneration activities in *nrl*-KO zebrafish at 9 and 18 mpf when compared with WT controls. Scale bars: 50 μm. (F) Quantitative results of the Pcna^+^ cells located in ONL per section are shown as mean with SD (n=4) from 7 dpf to 29 mpf. *, p < 0.05; **, p < 0.01.

Interestingly, we found that the retinal degeneration progression in *nrl*-KO zebrafish is quite slow according to our histological and immunofluorescence results, in spite of the evident apoptosis of retinal cells. Progressive photoreceptor degeneration has been reported to activate regeneration in zebrafish[50]. We speculated that retinal regeneration might also be activated to replenish the lost rods in *nrl*-KO zebrafish. Immunostaining of Pcna on retinal sections indeed showed more proliferating cells in *nrl*-KO retinas since 9 mpf (Fig 7E and 7f). These results suggested that progressive retinal degeneration in *nrl*-KO zebrafish might induce regeneration of photoreceptors in an *nrl*-independent manner.

## Discussion

Zebrafish has becoming a popular animal model for retinal degeneration and regeneration studies [24-26] and drug screening [51,52]. The developmental models of photoreceptors in zebrafish are established mainly based on the assumption that the functions of key regulatory genes (such as *NRL* and *NR2E3*) are conserved between zebrafish and mammals. However, this assumption has been challenged by several recent studies [29-31]. In this study, we generated an *nrl* knockout zebrafish model and investigated the development and degeneration of photoreceptors. We also determined the retinal cell populations and the cell-type-specific gene expression profiles in both WT and *nrl* knockout zebrafish through single-cell RNA-seq. The unexpected retinal phenotypes and the cellular and molecular findings described here expanded the current developmental model of rod photoreceptors in zebrafish (Fig 8).

**Fig 8.**
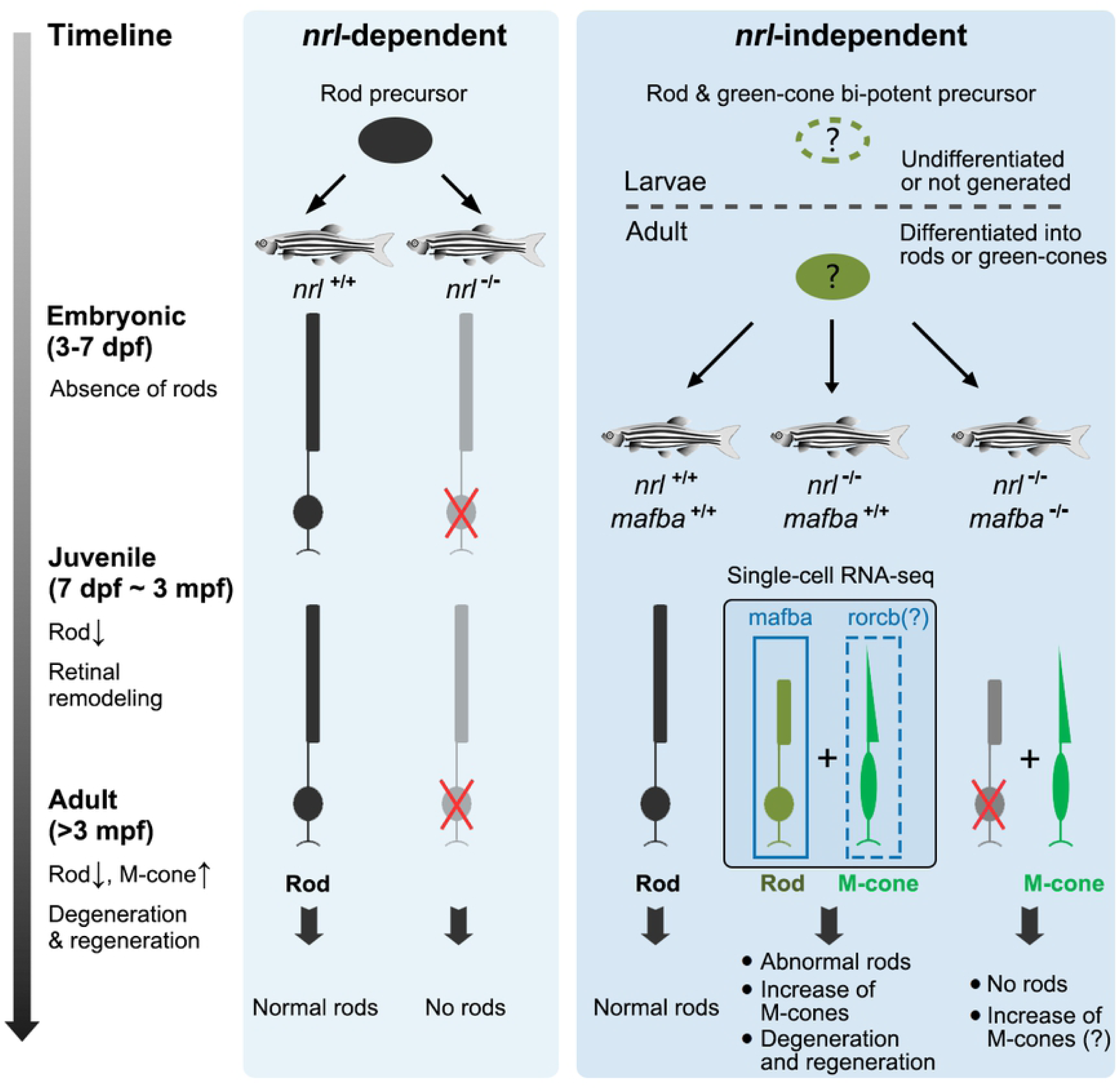
The supposed developmental model of rod photoreceptors in zebrafish. There may be two types of rod precursors defined as *nrl*-dependent and *nrl*-independent in zebrafish retinas. The former are responsible for the genesis of rods at embryonic stage, and knockout of *nrl* abolishes their differentiation into rods (left panel). The latter are responsible for the genesis of rods at the juvenile and adult stages. In the presence of *nrl* and *mafba*, these cells differentiate into rods. Knockout of *nrl* solely doesn’t severely affect the production of rods from these cells, but causes the abnormal expression of rod- and cone-specific genes and the gradual increase of green-cones (M-cones). Double knockout of *nrl* and *mafba* eliminates all rods, suggesting that *mafba* is the fate-determining gene for this type of rods (right panel).

Interestingly, we found that some features of *nrl* knockout zebrafish are more similar to the *rd7* mice carrying a *Nr2e3* mutation [32], such as the moderately decreased expression of rod-specific genes and the rod-cone “hybrid” cells (Fig 3-6). Meanwhile, the *nr2e3* knockout zebrafish show no rods and also no expression of rod-specific genes [30], which is more similar to the situation observed in *Nrl* knockout mice [10]. Moreover, *Nr2e3* is a direct target gene of *Nrl* in humans and mice [23]. However, knockout of *nrl* doesn’t significantly affect the expression of *nr2e3* in zebrafish (Fig 6C). A previous study has shown that *nr2e3* is expressed prior to *nrl* during rod genesis in zebrafish [28]. Taken together, we speculate that *nrl* might act in parallel with or even downstream of *nr2e3* in zebrafish. The functions and regulatory relationship of *NRL* and *NR2E3* are not totally conserved between fishes and mammals.

The gradual loss of rods results in secondary death of cones in retinitis pigmentosa, the most common type of inherited retinal degenerative disease [48,49]. The reduction of rods in *nrl* knockout retinas could mimic this situation and cause progressive retinal degeneration, as validated by the elevated levels of apoptosis of retinal cells and significant up-regulation of GFAP in Muller glia (Fig 7). In addition, our bulk and single-cell RNA-seq data may also provide valuable clues for investigating the genes and pathways involved in the occurrence and progression of retinal degeneration. For example, several differentially expressed genes, including *aipl1* (aryl hydrocarbon receptor interacting protein-like 1), *ahrrb* (aryl-hydrocarbon receptor repressor b), *rpgra* (retinitis pigmentosa GTPase regulator a), *nxnl1* (nucleoredoxin like 1), have been associated with retinal degenerative diseases previously [53-57]. Our study suggested that the *nrl* knockout zebrafish may be a novel and excellent model for investigating the molecular mechanism underlying the death of photoreceptors.

One of the major findings in this study is the discovery of two waves of rod genesis emerging at the embryonic and post-embryonic stages with *nrl*-dependent and *nrl*-independent patterns, respectively. Based on the temporal and spatial features, we suspect that the second wave of *nrl*-independent rods may likely represent the rod population constantly generated from progenitors in the inner nuclear layer during retinal expansion and growth of zebrafish [28,58]. Interestingly, a recent study have also reported the dispensable role of *nrl* in the specification of rods in adult zebrafish but not zebrafish larvae [59], which is partly consistent with our conclusions. Through single-cell RNA-seq, we further demonstrated the existence of different rod subtypes (Fig 5) and revealed their signatures in terms of developmental processes and gene expression patterns. These findings explain the unique retinal phenotypes observed in *nrl* knockout zebrafish, and suggest that zebrafish rods may be derived from different precursors under the control of regulatory genes such as *nrl* and *mafba*.

The gradually increased green-cones and the rod/green-cone “hybrid” photoreceptors in *nrl* knockout zebrafish suggest that there may be cell fate transformation between rods and green-cones. Opsins of rods and green-cones indeed show the highest similarity and the closest evolutionary relationship in non-mammalian vertebrates [60]. We speculated that at least a proportion of rods and green-cones may originate from the same precursors in zebrafish, and *nrl* may play a crucial role in the fate choice between rods and green-cones. However, the RH2 (green-cone) and SWS2 opsins are thought to be lost in mammalian ancestors during the adaption of nocturnal lifestyle [61]. This may explain why the S-cones but not the green-cones are alternatively increased in *Nrl* and *Nr2e3* knockout mice [10,11,32,37].

Recently, scRNA-seq has emerged as a powerful tool to study retinal development and degeneration under normal or pathological conditions [62]. In this study, the cell compositions and cell-type-specific gene expression profiles in WT and *nrl* knockout retinas were identified by scRNA-seq. Using these valuable data, we successfully identified the *mafba* gene as novel driving factor for the *nrl*-independent rods, which was subsequently validated by the *nrl* and *mafba* double knockout zebrafish model. Moreover, we identified a cone-specific nuclear receptor gene *rorcb* (RAR-related orphan receptor C b) that is up-regulated in *nrl* knockout retinas (Fig 3A and 6A). Whether *rorcb* is responsible for the overgrowth of green-cones in *nrl* knockout zebrafish needs to be determined in future studies.

In summary, we comprehensively identified the retinal phenotypes of *nrl* knockout zebrafish at cellular and molecular levels, and discovered novel roles of *nrl* and *mafba* in rod genesis, which contributes to a more suitable developmental model of photoreceptors in zebrafish. Our findings may promote further studies in the fields of photoreceptor development and evolution, and retinal degeneration and regeneration.

## Materials and methods

### Zebrafish lines

Wild-type (WT), *nrl* and *mafba* mutant, and Tg(rho:EGFP) transgene zebrafish were maintained and bred following the previous study [63]. The *nrl* and *mafba* knockout zebrafish were generated by CRISPR-Cas9 technology as previously described [30,64]. Zebrafish were sacrificed by immersing in 0.02% MS222 (Sigma, Cat# 886-86-2) solution for 15 minutes until no opercular movement. All procedures involving zebrafish were approved by the Ethics Committee of Huazhong University of Science and Technology.

### Retinal sectioning and HE (Hematoxylin and Eosin) staining

Zebrafish eyes were isolated and fixed with 4% PFA in PBS at 4°C overnight. After cryo-protection in 30% sucrose at 4°C overnight, the eyes were embedded in OCT (SAKURA) and sectioned along with the dorsal-ventral orientation through the optic nerve, using a cryostat microtome (CM1860, Leica). The thickness of the slices was set to 10-15 μm. The slides were dried at 37°C for 30 minutes and then stored at -20°C. HE staining was performed according to the instructions. The slides were observed and photographed under a light microscope (BX53, Olympus).

### Western blot and qPCR

Zebrafish eyes were enucleated for protein and RNA extractions. For each protein sample, 2-3 eyes from different zebrafish were sonicated in RIPA lysis buffer containing protease inhibitor cocktail (Sigma, Cat# P2714). Protein samples were mixed with loading buffer and boiled for 5 minutes at 95-100°C, and then stored at -20°C. For each RNA sample, three eyes (lens removed) from different zebrafish were homogenized in RNAiso Plus reagent (Takara, Cat# 9108). Total RNAs were extracted following the operation manual and stored at -70°C. Western blot and qPCR were performed as described previously [63]. The antibodies and primers used in this study are listed in S1 Table and S2 Table, respectively.

### Immunofluorescence staining and TUNEL assay

Immunofluorescence staining on retinal sections was performed as described previously [63]. For retinal whole-mount preparation, zebrafish eyes were fixed in 4% PFA in PBS at room temperature for 30 minutes and dissected under a stereomicroscope. The anterior segment, sclera, and choroid were removed, and the retinas were cut into several pieces. The whole-mount retinas were re-fixed in 4% PFA for 30 minutes, washed 3 times in PBS, blocked and permeabilized in PBST (0.5% Triton X-100) + 10% goat serum overnight at 4°C. The retinas were incubated with the primary antibody solution overnight at 4°C, washed 3 times in PBST for 30 minutes each, incubated with the secondary antibody solution for 4 hours at room temperature, and washed another 3 times in PBST for 30 minutes. To show the entire retina, a series of photographs showing the different regions of the same retina were combined into a panoramic image. TUNEL (terminal deoxynucleotidyl transferase biotin-dUTP nick end labeling) assay was performed on retinal sections following the manuals (Cat # 11684795910, Roche).

### Tracing the fluorescence of rods in Tg(rho:EGFP) transgene zebrafish

At 3-17 dpf, the fish were sacrificed and fixed in 4% PFA in PBS for 15 minutes at room temperature. After being washed three times in PBS, the eyeballs were isolated and the sclera/choroid was removed under a stereomicroscope. The eyeballs were depigmented in 3% H_2_O_2_ + 1% KOH solution at room temperature for 5-10 minutes, washed 3 times in PBS for 10 minutes each, mounted in 0.8% low-melting agarose, and photographed immediately under the fluorescence microscope (Eclipse 80i, Nikon). To show the fluorescence of the whole retina more comprehensively, two photographs with different Z-positions were taken and merged into one image for each eyeball.

At 1-3 mpf, the zebrafish were sacrificed and the eyeballs were isolated and fixed in 4% PFA for 30 minutes at room temperature. After being washed three times in PBS, the anterior segment, sclera, and choroid were removed under a stereomicroscope. The retinas were depigmented in 3% H_2_O_2_ + 1% KOH solution at room temperature for 5-10 minutes, washed 3 times in PBS for 10 minutes each, cut into several pieces, and mounted in PBS on microslides. Photographs were taken immediately under the fluorescence microscope (Eclipse 80i, Nikon). A series of local images from a single retina were merged to show the whole view.

### *In situ* hybridization on retinal sections

The cDNA fragment of *hmgn2* was amplified from the retina cDNA library by primers (F: CGCGAGGTTGTCTGCTAAAC, R: GGTGAAAACCCTTCCGAAAACA) and validated by Sanger sequencing. The T7 promoter was added to the 5’ and 3’ ends to synthesize the sense and anti-sense probes using the DIG RNA Labeling Kit (Roche, Cat# 11175025910). *In situ* hybridization was performed as previously described [65,66] with minor modifications. The signals were detected and developed by the Anti-Digoxigenin-AP antibody (Roche, Cat# 11093274910) and NBT/BCIP solution (Roche, Cat# 11681451001) following the manuals. The slides were examined and photographed under a light microscope (BX53, Olympus).

### RNA-seq and bioinformatic analysis

The total RNA samples were quantified and qualified by Agilent 2100 Bioanalyzer (Agilent Technologies) and NanoDrop (Thermo Fisher Scientific). RNA samples with RIN ≥9.5 and A260/280 ≥1.9 were used for RNA-seq. The next-generation sequencing and data analysis were performed by GENEWIZ (Suzhou, China). Briefly, the NEBNext Ultra RNA Library Prep Kit for Illumina was used for library preparations. Next-generation sequencing was carried out on the Illumina HiSeq instrument using a 2×150 bp paired-end (PE) configuration according to the manufacturer’s protocol.

The high-quality clean data were generated by Trimmomatic (v0.30) and aligned to the reference genome of zebrafish (GRCz11 from Ensembl) via software Hisat2 (v2.0.1). Gene and isoform expression levels were estimated by HTSeq (v0.6.1). Differential expression analysis was performed by DESeq Bioconductor package. After adjustment by Benjamini and Hochberg’s approach for controlling the false discovery rate, the threshold of p-values was set to <0.05 to detect differentially expressed genes. GO and KEGG enrichment analysis was then performed to enrich differential expression genes in GO terms and KEGG pathways.

### Preparation of the retinal single-cell suspension

The eyes of 5-month-old WT and nrl-KO zebrafish were isolated and dissected in cold HBSS buffer using microsurgical scissors and forceps under a stereomicroscope. The anterior segment, sclera, and choroid were removed. The retina was transferred to a well of a 12-well plate with 1 ml of the activated papain dissociation solution (20 U/ml papain, 1 mM L-cysteine, 0.5 mM EDTA in HBSS, pH 6.0-7.0; activated by incubation at room temperature for 30 minutes), and incubated at 28°C for 10-15 minutes with gentle agitation. The retina was aspirated and expelled through a wide orifice pipette tip every 2 minutes to promote dissociation and prevent aggregation. The visible pieces of non-dissociated tissues were removed with fine forceps. The cell suspension was filtrated through a 40 µm cell strainer (Corning, Cat# 431750), and then pelleted through centrifugation at 300g for 5 minutes at room temperature. The cells were washed twice in HBSS buffer (no Ca^2+^ and Mg^2+^) with 0.04% BSA and resuspended in the same buffer. Cell viability was determined by trypan blue staining, and should be over 80%. Cell concentration was adjusted to ∼1000 cells/ul. The cell suspension was placed on ice (<30 minutes) and ready for scRNA-seq.

### Single-cell RNA-seq (scRNA-seq) and bioinformatic analysis

Retinal single-cell suspensions were loaded onto the 10x Genomics Chromium Single Cell system using the 3’ Reagent Kits (v3 chemistry) at Genedenovo Biological Technology Co., Ltd. (Guangzhou, China). Approximately 10,000 live cells were loaded per sample. Single-cell GEM generation, barcoding, and libraries preparation were performed according to the user guide. High-throughput sequencing was performed on the HiSeq 2500 systems (Illumina) at Genedenovo. The data were analyzed using Cell Ranger pipelines to align reads to the reference genome (GRCz11 from Ensembl), generate feature-barcode matrices, and perform clustering and gene expression analysis.

Seurat <u>_ENREF_57</u>[67] was used for quality control, gene expression normalization, principal component analysis (PCA), and cell clustering. In brief, cells with a clear outlier number of genes (potential multiplets) and a high percentage of mitochondrial genes (potential dead cells) were excluded by “FilterCells” function. A global-scaling normalization method “LogNormalize” was employed to normalize the gene expression measurements for each cell. PCA was performed to reduce dimensionality of the dataset. Seurat clustered cells based on their PCA scores. To determine the most contributing PCs, a resampling test inspired by the jackStraw procedure was implemented. The significant PCs with a strong enrichment of low p-value genes were used to identify the unsupervised cell clusters. t-SNE (t-distributed Stochastic Neighbor Embedding) was used to visualize and explore the distribution of cells in a low-dimensional space (2D-plot). For every single cluster, differentially expressed genes (up-regulated) were identified by the likelihood-ratio test using “FindAllMarkers” function in Seurat package, compared to all other cells. The top 20 differentially expressed genes were selected as marker genes for each cluster. GO and KEGG enrichment analysis was performed to recognize the main biological functions and the significantly enriched metabolic or signal transduction pathways in differentially expressed genes.

### Statistical analysis

Unpaired Student’s two-tailed t-test was performed to determine the statistical significance between two groups in GraphPad Prism 6 software. A p-value less than 0.05 was considered statistically significant (*, p < 0.05; **, p < 0.01; ***, p < 0.001). All data are presented as mean with SD. All experiments were repeated at least three times independently.

## Data availability

The NGS data are available under the GEO accession numbers: GSE160138 (RNA-seq) and GSE160140 (scRNA-seq). Other relevant data are within the manuscript and its Supporting Information files.

## Acknowledgments

We thank Dr. Jian Zou (Eye Center of the Second Affiliated Hospital School of Medicine, Institutes of Translational Medicine, Zhejiang University) for providing the Tg(rho:EGFP) zebrafish line. This work was supported by the National Natural Science Foundation of China [No.31601026, No.81670890, No.31801041, and No.31871260] and the National key R&D Program of China [No. 2018YFA0801000].

## Author Contributions

Conceptualization: M.L., F.L., D.L., C.X.; Data curation: M.L., F.L.; Formal Analysis: F.L., Y.Q.; Funding acquisition: M.L., Z.T., C.X., F.L., S.Y.; Investigation: F.L., Y.Q., Y.H., P.G.; Methodology: F.L., Y.Q., Y.H., P.G., X.Z., J.L.; Project administration: M.L., Z.T., F.L.; Resources: K.S., D.J., X.C., Y.L., Y.H., J.T.; Supervision: M.L., F.L., D.L.; Visualization: F.L., Y.Q.; Writing – original draft: F.L., Y.Q.; Writing – review & editing: M.L., F.L., D.L., C.X., Q.L., X.S., J.R..

## Declaration of Interests

The authors declare no competing interests.

## Notes

### Competing Interest Statement

The authors have declared no competing interest.

